# Enhancer-promoter association determines *Sox2* transcription regulation in mouse pluripotent cells

**DOI:** 10.1101/590745

**Authors:** Lei Huang, Qing Li, Qitong Huang, Siyuan Kong, Xiusheng Zhu, Yanling Peng, Yubo Zhang

## Abstract

Distal enhancer-promoter associations are essential for transcription control and emerging as a common epigenetic way to determine gene expression in eukaryotes. However, it remains as an uncultivated land on how their diversity influences gene transcription during development. In this study, we select three defined *Sox2* associated enhancers (E1, E2 and E3), representing different distal interaction categories, to further explore their biological effects. We construct three enhancer-knockout cells via CRISPR/Cas9 system in mouse embryonic stem cells. Results show that these associations carry out various biological features. Embryonic stem cell specific association (E1) speeds up cell cycle and involves in cardiac development, which has been validated *in vivo.* In contrast, the indirect one (E2) restrains nerve differentiation and has potential effect on lipid metabolism. The common one (E3) promotes nerve differentiation and inhibits oxidation-reduction process. Together, these different associations determine *Sox2* transcription and function specificity in mouse embryonic stem cells. Our study will enable a way for exploring miscellaneous spatiotemporal gene transcription control and advancing quantitative knowledge by utilizing three-dimensional genomic information.

**Summary statement:** Sox2 transcription is complicated in mouse embryonic stem cells. It involves in heavily enhancer-promoter associations, which might make it a suitable model for phase-separation study. To decompose this cooperative regulation, diverse interactions are investigated. The biological effects have been explored in multiple perspectives. This may represent a quantitative way to explore and a potential new strategy to transcription control study.

## Introduction

Mammalian development and cell identity heavily depend on quantitative transcription control to establish spatiotemporal gene expression programs (Hnisz et al., 2017; Rubin et al., 2017; Weintraub et al., 2017). Recent genome-wide analyses have revealed that long range enhancers positively regulate transcription through physical contacts with gene promoters during their activation (Li et al., 2012a; DeMare et al., 2013; Kieffer-Kwon et al., 2013; Dowen et al., 2014; Ji et al., 2016; Liu et al., 2017). Enhancer activity involves in diverse interaction with coactivators and promoter, which accompany the formation of transcription complex and has potential influence on gene regulation network (Kleinjan and van Heyningen, 2005; Rubtsov et al., 2006; Jager et al., 2015). Recently, a phase separation model is proposed as a general regulatory mechanism to compartmentalize biochemical reactions within cells (Hnisz et al., 2017). Prominent roles of enhancers on activating genes with diverse enhancer-promoter (E-P) associations are proposed as well. Developmental genes are often controlled by multiple enhancers, which can cooperate to induce transcription of their shared target gene (Krijger and de Laat, 2016; Long et al., 2016). Recently, a research implies that enhancer RNAs may serve as bridge for cooperation of multiple enhancers(Tsai et al., 2018). However, the regulatory relationships among simultaneously active enhancers are still unclear. To quantitative or precisely measure gene transcription, it will be essential to decompose their biological effects.

*Sox2* is a key transcription factor (TF) for maintaining pluripotency and self-renewal in embryonic stem cells (ESCs) (Masui et al., 2007). The transcription regulation of *Sox2* gene is complicated in ESCs. On one hand, its function highly depends on cooperating gene expression with other TFs such as *Oct4, Nanog, Esrrb, Tbx3,* and *Tcf3*(Ang et al., 2011; Narva et al., 2012). On the other hand, E-P association has been reported to be dispensable during activation (Masui et al., 2007). For example, *Sox2* enhancers can form 3D-clusters which are overlap with a subset of Pol II enriched regions (Liu et al., 2014). Meanwhile, in mouse ESCs (mESCs), the highly expressed *Sox2* can be repressed by P21 or P27 which target two proximal enhancers and recruit transcription inhibitors (Sikorska et al., 2008; Li et al., 2012b; Yamamizu et al., 2014). A distal downstream enhancer has been reported that it could tightly interact with *Sox2* promoter and is essential for appropriate transcription in mESCs (Li et al., 2014; Zhou et al., 2014). Depletion or epigenetic modification of these enhancers will attenuate the pluripotency of mESCs(Li et al., 2014; Zhou et al., 2014). However, the cooperative biological effect of different E-P associations of *Sox2* gene has seldom been explored.

In this study, we have identified three distal enhancers (termed E1, E2, E3), interacted with *Sox2* promoter in different manners (Zhang et al., 2013). They represent specific (E1), indirect (E2) and common (E3) E-P associations with *Sox2* gene in mESCs. Among them, E1 forms specific interaction with *Sox2,* E3 is a common interaction which could be found both in mESCs and mouse neural stem cells (mNSCs). E2 is a known enhancer in mouse neural progenitor cells (mNPCs) and has no direct interaction in mESCs with current datasets (Figure 1A). To further identify the diverse biological effect, we construct three enhancer knockout cells via CRISPR/Cas9 system in mESCs. Together with genomic and molecular evidences, we have explored their various biological effects and uncover potential pattern of *Sox2* gene transcription control associated with E-P interactions.

**Figure 1.**
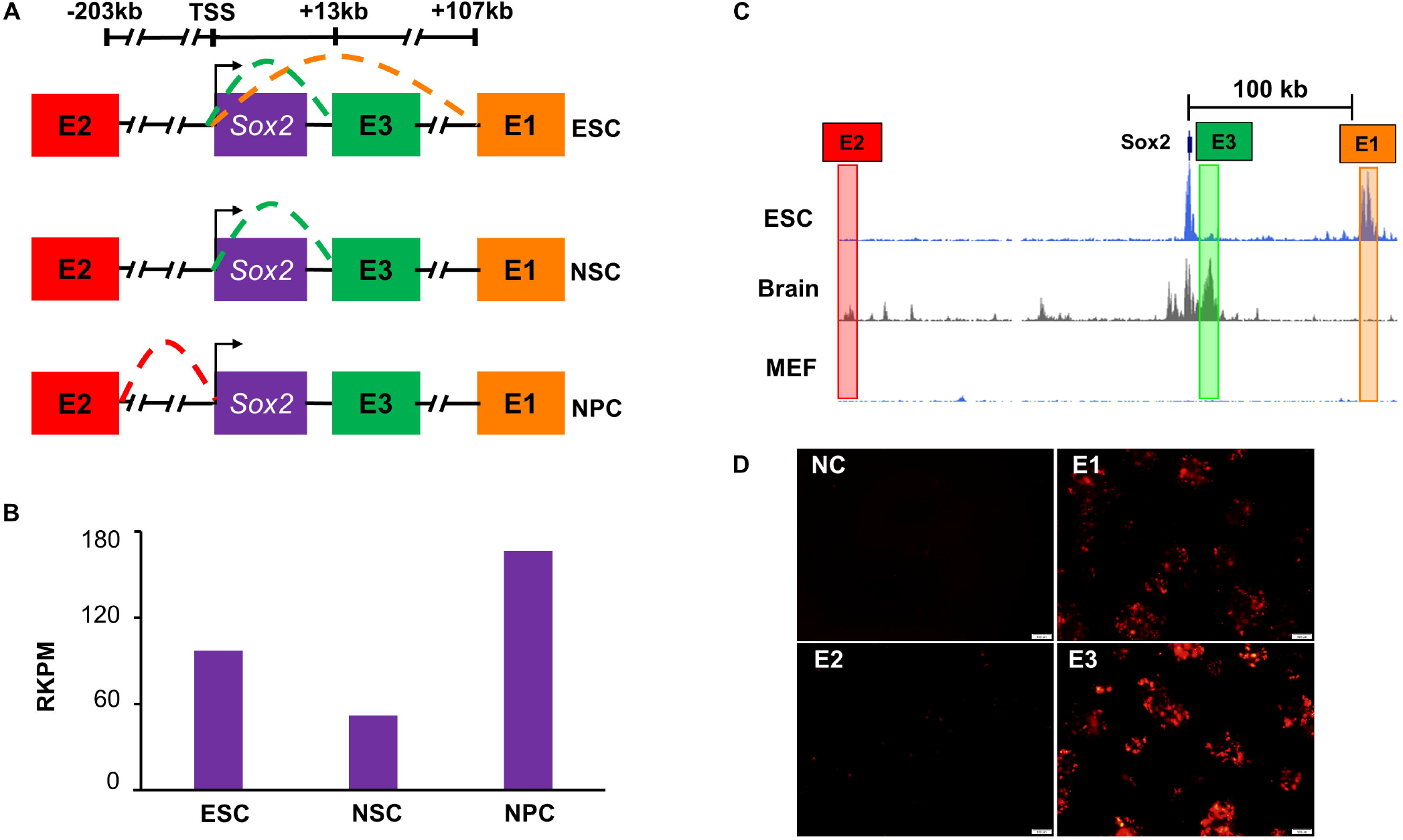
Different patterns of E-P association and activity assay of *Sox2* enhancer. (A) Three enhancers of *Sox2* gene exhibited different patterns of E-P association. The above line shows the distance from TSS. The three *Sox2* enhancers named E1, E2, and E3 demonstrated with different colored block (orange, red and green). Patterns of E-P association are denoted with corresponding colored dotted line. (B) The expression level of *Sox2* genes in mESCs, NSC and NPC. The RKPM value derived from RNA-seq data of three cell lines. (C) ChIP-seq peaks for H3K27ac in mESCs, brain and mouse embryonic fibroblast (MEF). The region of E1, E2 and E3 are displayed in orange, red and green color respectively. (D) Enhancer activity assay. The enhancer activity is indicated with red fluorescence depending on pGL4.23-mCherry vector. NC, empty pGL4.23-mCherry vector; E1, E2, E3, inserted with corresponding enhancer fragment.

## Results

### Diverse E-P associations of *Sox2* gene

In our study, we validate the enhancer activity with histone marker and fluorescence labeling. Results indicate that, specific (E1) and common (E3) interaction show significant enhancer activity, while indirect (E2) interaction shows weak enhancer activity in mESCs (Figure 1C, 1D). Similar results are observed in 293T cells (Figure S1A) not in MEF (No obvious signals are found). In previous study, *Sox2* knockout mouse/cell leads to early mortality after implantation(Avilion et al., 2003; Adachi et al., 2013). Scientist can only generate conditional knockout mice to study *Sox2* gene function(Avilion et al., 2003). Additionally, *Sox2* is highly expressed in multiple mouse cells (Figure 1B), which makes obstacles for its function analysis. To avoid this shortcoming and further explore their transcription control, we construct three enhancer knockout (E-KO) cells via CRISPR/Cas9 system with mESCs (Figure 2A, S2A). All cells display normal clone morphology like wild type (WT) mESCs except E3-KO cells (Figure 2A, Figure S9C). At the same time, qPCR detection reveals that deletion of these three enhancers significantly reduces the expression of *Sox2* gene (*p*-value < 0.01, t-test) (Figure 2B). Therefore, these E-P associations of *Sox2* gene can affect its expression in both common and specific way (Figure S12).

**Figure 2.**
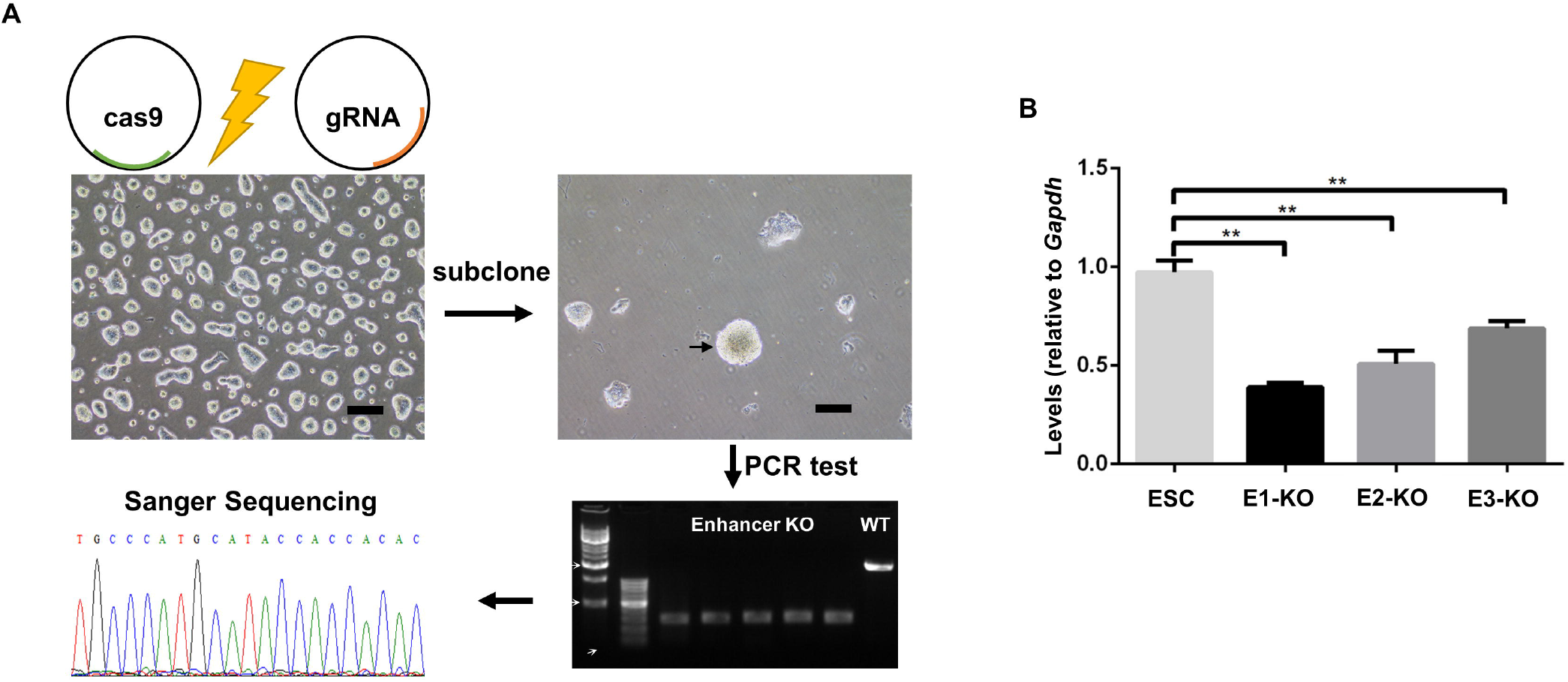
Deletion of enhancers in mESCs using CRISPR/Cas9. (A) The experimental process of enhancers knockout, including cell electro transfection, monoclonal selection, genotype identification and sanger sequencing. The ruler of the mESCs cell figure is 200 μm. Black arrow indicates the clone to be picked. (B) Expression of *Sox2* gene is detected in E-KO and WT mESCs cells. Values are an average of three biological replicate experiments, with each experiment performed in triplicate. Error bars represent SEM. A significantly lower expression level over the WT mESCs is indicated by double asterisks (*P* < 0.01, t test).

### RNA-seq datasets reveal E-P association deletion affecting *Sox2* gene transcription

Transcriptome assay is carried out using these E-KO cells (two biological replicates in each group). Differentially expressed genes (DEGs) analysis reveals that deletion of enhancers affects the transcription level of hundreds of genes in mESCs (q-value<0.05, Benjamini-Hochberg correction for *p*-value (t-test), | log2 (fold change) | >1). Compared with WT mESCs, E1-KO has 1209 DEGs (697 down, 512 up), E2-KO has 1020 DEGs (585 down, 435 up), and E3-KO has 1494 DEGs (938 down, 556 up) (Figure 3A, 3B and S3A) (Table S2). The number of down-regulated genes is a bit more than that of up-regulated genes. Meanwhile, the biological replicates in the experimental tests have a high correlation (The square of the Pearson correlation coefficient is greater than 0.98) (Figure 3C), indicating the consistency of the RNA-seq data. However, the square of correlation coefficient among the E-KO and WT mESCs reaches more than 0.94, reveals that the knockout of the enhancer has a weak influence on the gene expression level. In addition, the number of DEGs that changes 2-4 times accounted for around 80% of the total DEGs (Figure S4).

**Figure 3.**
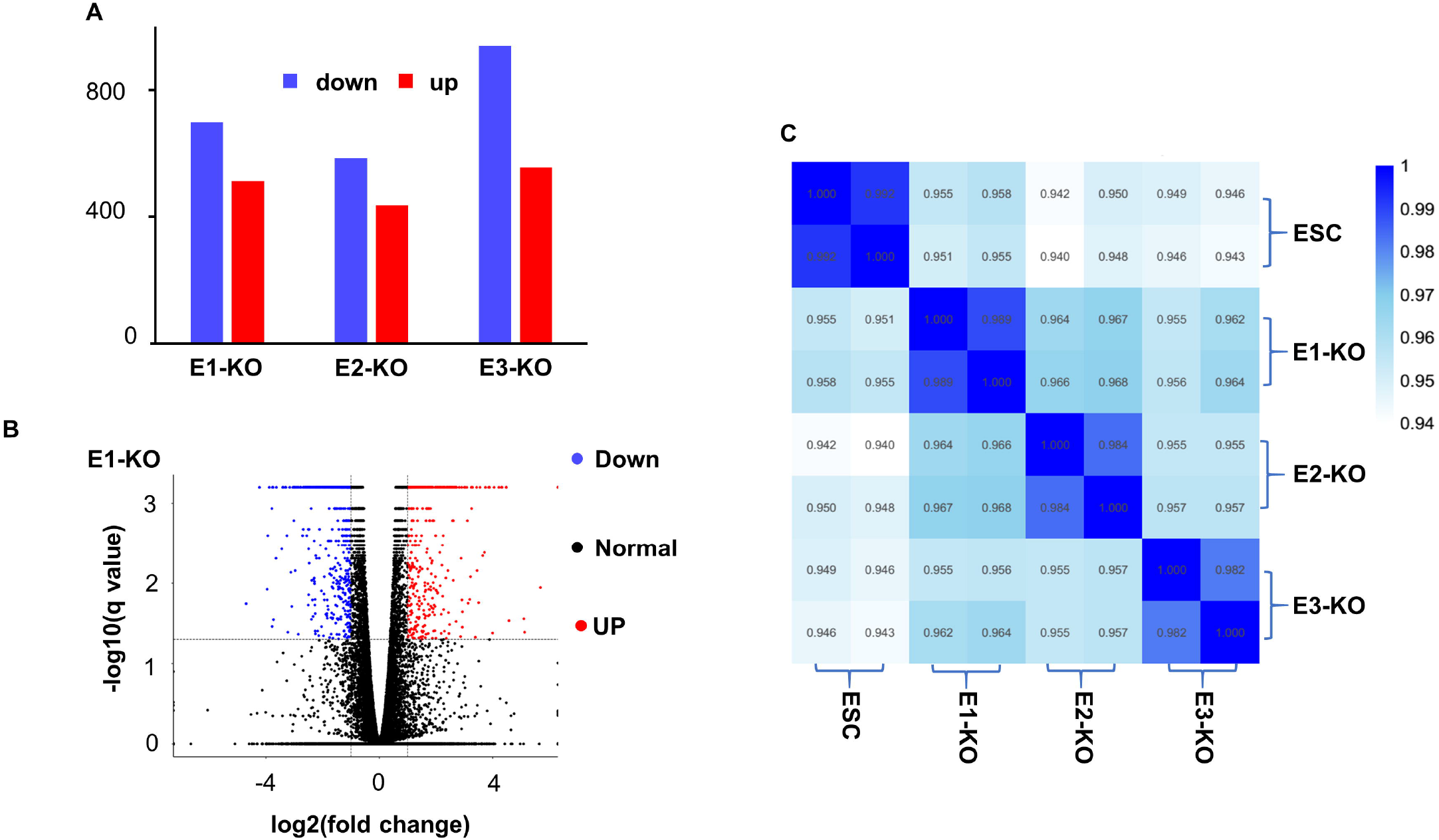
Analysis of differentially expressed genes derived from RNA-seq data of E-KO cells. (A) Statistics of the number of DEGs. E1-KO, E2-KO and E3-KO represent DEGs compared with WT mESCs. Blue bar, down-regulated genes; red bar, up-regulated genes. (B) Volcano plots reveal transcript abundance of E1-KO cell. The y-axis corresponds to the expression values of -log10(q-value), and the x-axis displays the log2 fold change values. The blue dots represent down-regulated genes; the red dots represent up-regulated genes; the black dots represent normal genes. (C) Pearson correlation between samples. Pearson correlation coefficients among all samples are calculated based on the values of log10(FPKM+1). Different colors represent the squared correlation coefficient (R^2^) which ranges from 0.94 to 1.

With these results, enhancers show quantitative regulation without causing dramatic changes in gene expression or phenotype faultiness unlike *Sox2* gene knock-out experiment (Drissen et al., 2010; Rowan et al., 2010; Seitan et al., 2013). This might represent a precise or alternative way to study gene function.

### Specific interaction (E1) influences cell cycle and differentiation

Specific interaction locates at distal downstream from *Sox2* and specifically interacts with its promoter in mESCs (Figure 1A). To explore its biological effects, we conduct enrichment analysis of biological processes and signal pathways of DEGs in E1-KO cells. Results show that specific interaction mainly affects biological processes including cell cycle and differentiation (Figure 4A, S5A) (Table S3). We examine the cell cycle of E1-KO cells to verify the role of E1 in cell proliferation using cytometry flow. Compared with the WT mESCs, the G1 phase of E1-KO cells increases significantly. While the S phase decreased significantly (*p*-value < 0.01, t-test) (Figure 4C, S8C, S10), indicating that E1 affects the expression of related genes during the cycle transition of mESCs from G1 to S phase. On the other hand, there is no significant difference in proliferation rate between WT mESCs and E1-KO cell (Figure 4D).

**Figure 4.**
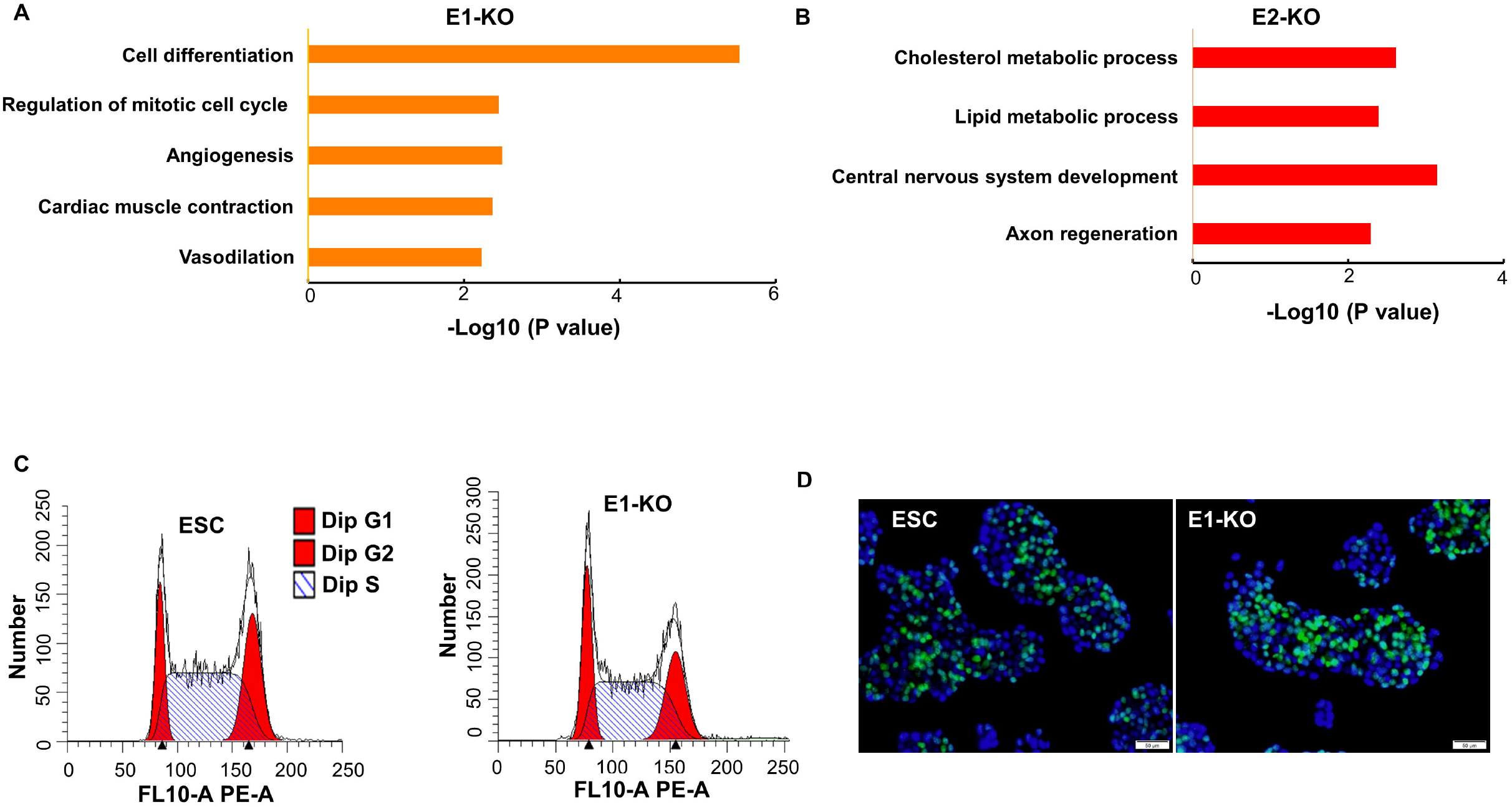
E-P interaction of E1 influenced proliferation, differentiation and cardiac function. (A, B) GO functional classification of DEGs from E1-KO and E2-KO cells. E1 is enriched in the process of cell differentiation, cell cycle and cardiac function, however, E2 is enriched in metabolic process and nervous development. The x-axis displays the values of -log10(*p* value). (C) Cell cycle distribution analyzed by the flow cytometry. The fitting period of G1, G2, S is achieved with Modifit software. (D) Cell proliferation assay with Edu dyeing. Blue, nuclear (represents total cells); green, anti-Edu (represents proliferating cells).

We also close observe the colony forming efficiency of different cells to characterize their feature effects on self-renewal. The result of colony formation assay reveals that the clonogenic rate of E1-KO cell is significantly lower than that of WT mESCs (*p*-value < 0.01, t-test) (Figure 5A and 5B). Meanwhile, DEGs KEGG pathway analysis also shows its enrichment in cell proliferation related pathways such as Ras, PI3K-Akt, and MAPK (Figure S5B) (Table S4). Results of hierarchical clustering of DEGs reveal subdivision of proliferation and differentiation related genes (Figure S8A) (Table S7). In addition, in the process of nerve differentiation, the expression of *Nestin* gene in E1-KO is significantly lower than WT mESCs (Figure S8B), indicating that specific interaction promotes nerve differentiation.

**Figure 5.**
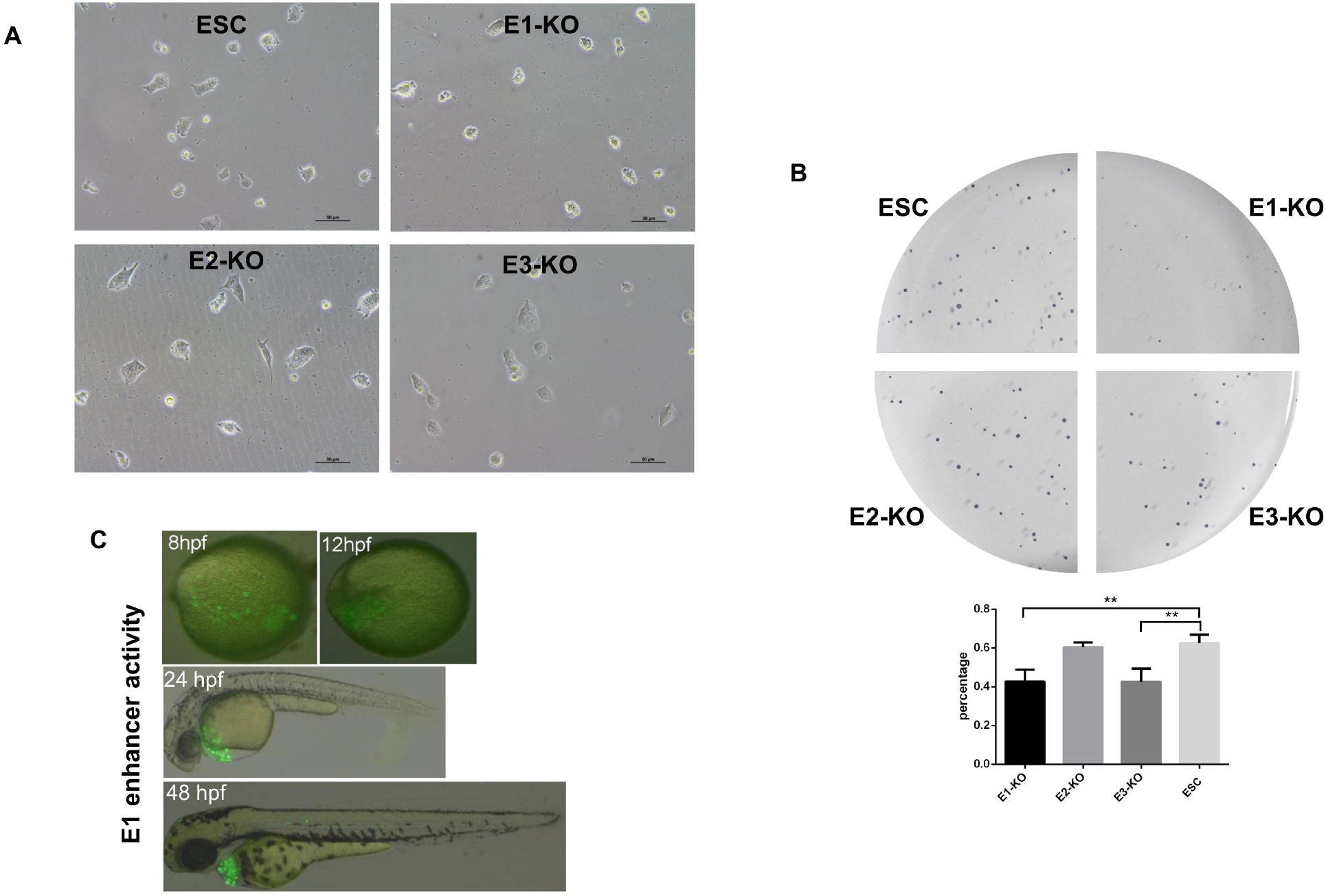
Enhancer E1 influences self-renewal of mESCs and displays activity in cardiac tissue of zebrafish. (A) The surviving cloned cells of E1-KO cells are significantly reduced after 24 hours of subculture. (B) The rate of clone formation of E1-KO cells decreases 7 days later. (C) E1 displays enhancer activity around heart of zebrafish in 24hpf and 48hpf.

Together, as an mESCs specific enhancer, specific interaction can maintain normal cell cycle and self-renew capability. At the same time, it has a positive effect on the differentiation in mESCs.

### Specific interaction (E1) determines in cardiac organ regulation *in vivo*

With further GO analysis results, we find a few genes associated with cardiac function in the DEGs of E1-KO cells. These are involved in the regulation of biological processes such as angiogenesis, cardiac muscle contraction, and vasodilation (Figure 6A, S5A) (Table S3). Pathway analysis also reveals that they enrich in pathway such as cardiomyopathy (Dilated, Hypertrophic, Arrhythmogenic) and cGMP-PKG (Figure S5B) (Table S4). Clustering analysis of cardiac function related genes shows specific subdivision of biological process (Figure 6A) (Table S5). Collectively, the up-regulated DEGs mainly promote angiogenesis. However, the down-regulated DEGs inhibit calcium ion transport and vasodilation. To validate *in vivo* regulation pattern, we construct the transgenic line in Zebrafish, which could investigate in transparent way during the early embryonic stages and offer an alternative method to validate vertebrate genes’ expression. In zebrafish, specific interaction appears enhancer activation at 24hpf in the heart primordium that migrate to the heart at 48hpf (Figure 5C, Figure S11). No signal is found in other organs. This indicates that specific interaction is involved in the development of the cardiovascular system and the regulation of cardiac function *in vivo*. Together, specific interaction can regulate gene function in cardiac during early embryonic period, which is consistent with our GO analysis.

**Figure 6.**
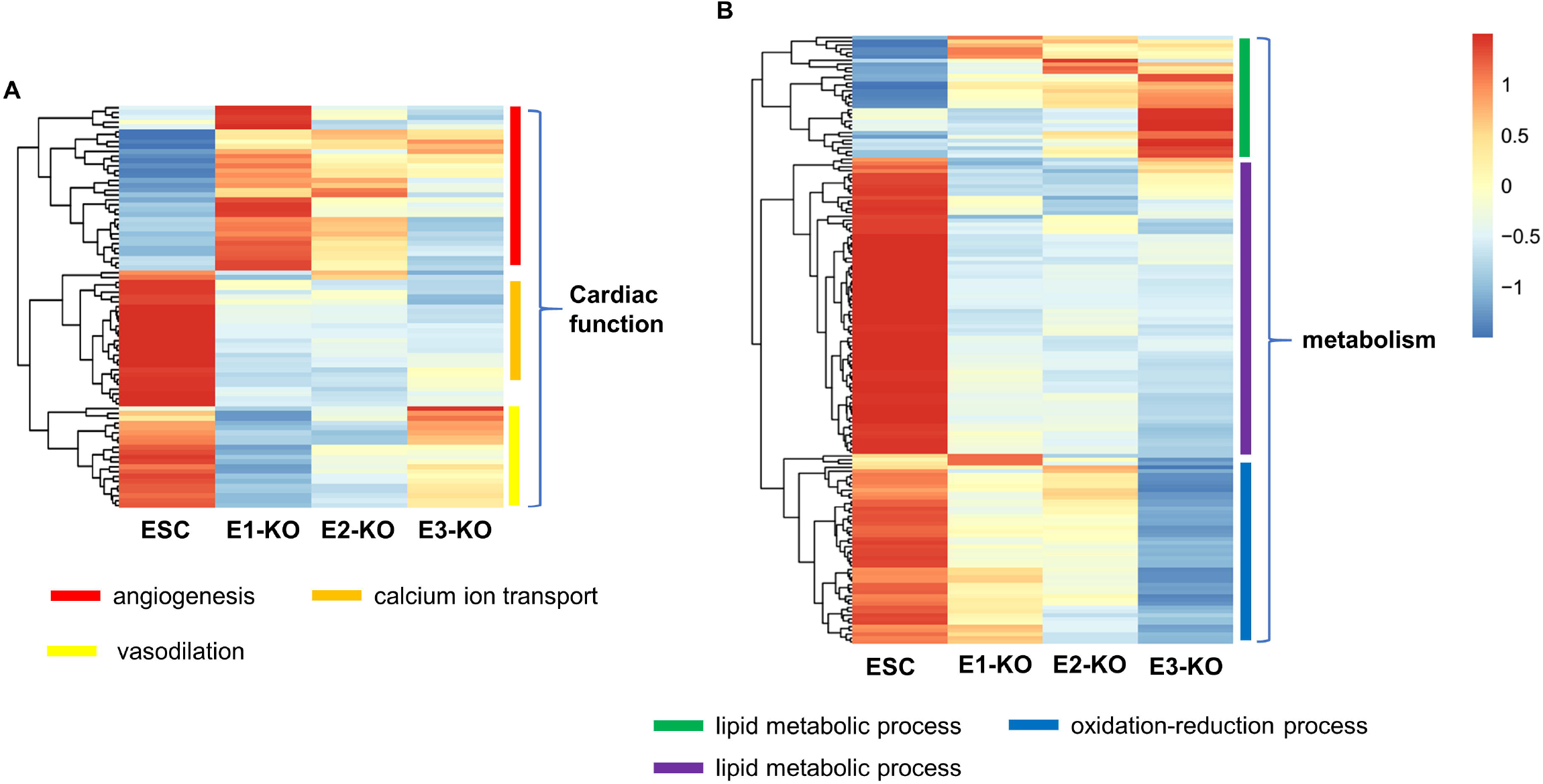
Gene expression heatmap of genes involved in cardiac function and metabolism. (A) Gene expression heatmap of genes involved in cardiac function. (B) Gene expression heatmap of genes involved in metabolism. Different colors represent the values of FPKM after z score transformation, (z scores are calculated by subtracting the overall average gene intensity from the raw intensity data for each gene and dividing that result by the SD of all the measured intensities). The clustering method is the complete link method in hierarchical clustering.

### Indirect interaction (E2) involves in metabolism and nervous system functions

Indirect interaction element cannot form any association with *Sox2* gene in mESCs. As a distal regulatory element upstream from *Sox2,* indirect interaction shows very weak enhancer activity in mESCs(Zhang et al., 2013). It lacks the histone markers of the active enhancers (Figure 1B). Hundreds of DEGs are found in RNA-seq data of E2-KO cells comparing with mESCs, indicating that indirect interaction could partially participate in transcription control in mESCs (Figure 3A and S3A). GO and KEGG analysis show that indirect interaction mainly relates to metabolism, cytoskeleton and nervous system functions (Figure 4B, S6A and S6B) (Table S3, S4). Indirect interaction involves in metabolic regulation including cholesterol, lipid, reactive oxygen, amino acid, glycan, *et al*. In addition, indirect interaction shows enhancer activity in nervous system, suggesting its potential regulatory role in nervous system function. GO analysis reveals those enrichment biological processes in nervous system development, neuron apoptotic, and axon regeneration confirmed such hypothesis (Table S3). Meanwhile, the expression of *Nestin* is significantly higher than that of WT mESCs on the fifth day of nerve differentiation (Figure S8B). These results provide *in vitro* evidence for the inhibition effect of indirect interaction in nervous system development.

Collectively, indirect interaction shows very weak enhancer activity and don’t have directly association with *Sox2* in mESCs. However, it could partially participate in the transcription control of other genes which could also affect some biological function of mESCs.

### Common interaction (E3) functions in ion transport and metabolism genes

Common interaction which has been reported mESCs and mNSCs(Zhang et al., 2013). In mESCs, common interaction has enhancer activity and interaction with *Sox2,* indicating the positive transcriptional effects as specific interaction. GO analysis shows that common interaction influences ion transport, ion homeostasis and metabolism of amino acids which are also regulated by specific or indirect interaction (Figure S2A and S7A) (Table S3). However, pathway analysis indicates that common interaction has a unique regulatory role in glycolysis/gluconeogenesis and pyruvate metabolism (Figure S7B). Clustering analysis of metabolism related genes reveals that its DEGs are divided into three subdivisions (Figure 6B) (Table S6). Concisely, common interaction shows different pattern in the regulation of lipid metabolic process and oxidation-reduction process. Furthermore, common interaction has significant enrichment in signaling pathways regulating pluripotency of stem cell. (Figure S9A and S9B) (Table S8). The clone morphology of E3-KO cells is relatively looser than WT mESCs implying lower pluripotency (Figure S9C)(Ying et al., 2008).

Together, common interaction affects ion transport and metabolism regulation which are partially overlapped with specific or indirect interaction. Besides that, common interaction might play essential role in the pluripotent maintenance of mESCs as well.

### E-P associations cooperate *Sox2* transcription with different genes

*Sox2* is the key node of the core transcription regulatory network in mESCs(Zhang et al., 2013). To further investigate the cooperative pattern and biological effects of these three enhancers, we extract gene interaction information to re-depict network of *Sox2* and our expression datasets (Figure 7). We find that the interactions among these enhancers (specific, common and indirect) and *Sox2* gene have different patterns to perform their biological functions. The mESCs specific E-P association (E1) is involved in cell differentiation and heart development. Its interaction with *Sox2* significantly reduces its expression level and affects the expression of *Sox2ot* and *Hist1h2ag* genes, which has also been shown in our GO gene list. The indirect E-P association (E2) is involved in cell differentiation and nucleosome assembly and participates in the regulation of other genes, such as *Myc* and *Pvt1.* The common E-P association (E3) is participated in cell growth and pluripotency and can influence more gene expression through several common interactions. It affects the expression of *Klf4* through the action of *Neat1* and affects the expression of *Hist1h4b* or *Hist1h1a* through *Setd5* or *Trim8* (Figure 7).

**Figure 7.**
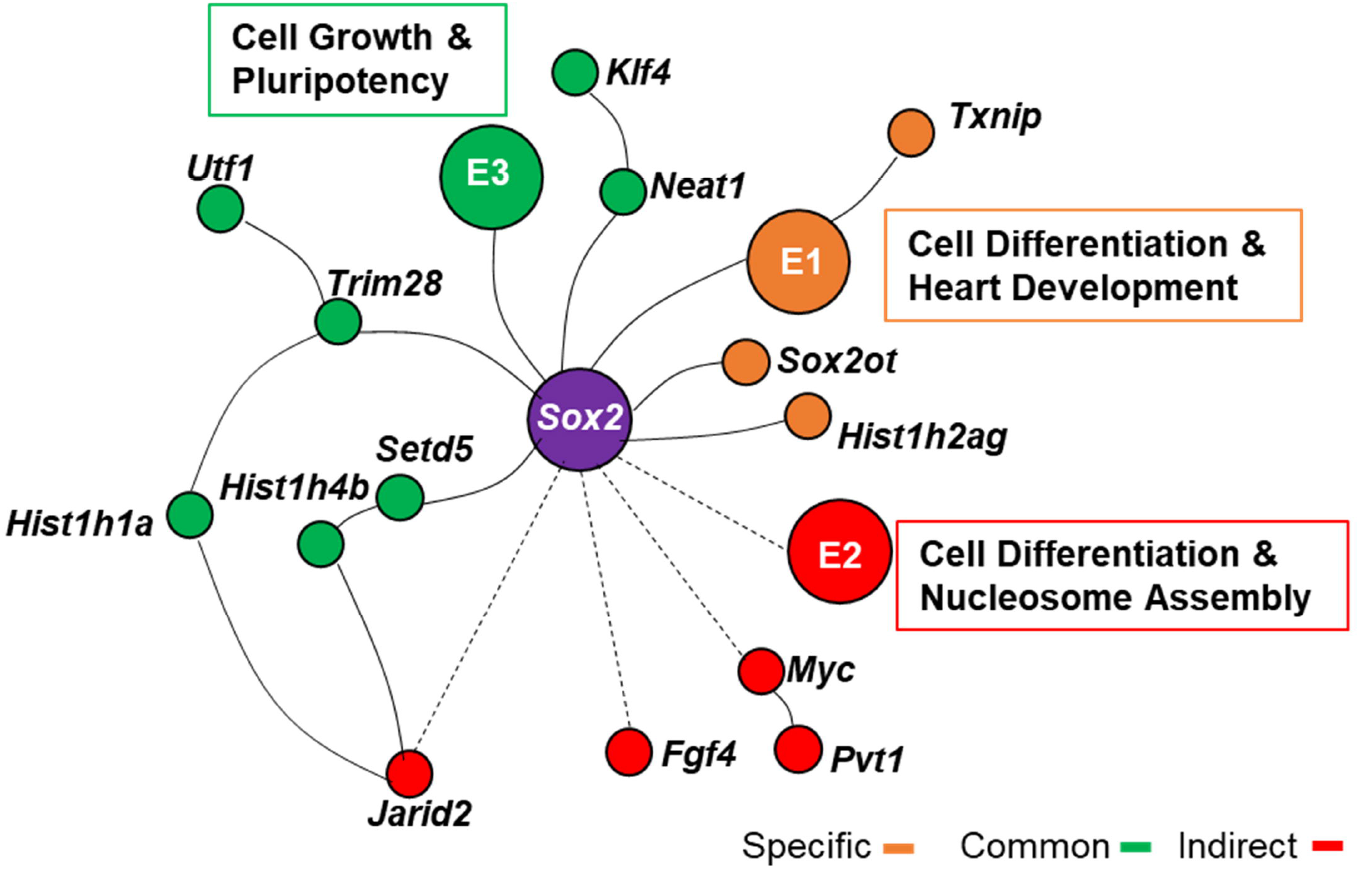
Gene network related with *Sox2.* Different patterns of interaction among genes connect with *Sox2* in direct or indirect way. Dotted line, indirect interaction; red circle, indirect genes; orange circle, specific genes; green circle, common genes.

Together, these imply diverse E-P associations could cooperate to control *Sox2* gene transcription with distinct individual features and determine different aspects of cell function. As a TF gene, the transcription control of *Sox2* behaves complicatedly and might fulfill its spatiotemporal regulation via distal interactions with multiple enhancers and genes in mESCs. And further effects are needed to decompose its diverse function.

## Discussion

*Sox2* gene transcription in mESCs is complicated and might be a good model for phase separation study. Up to now, it remains as challenge to study *Sox2* gene function with knockout system due to its importance in pluripotency and self-renew. Ablation of *Sox2* gene leads to mortality of mESCs and mice(Avilion et al., 2003; Arnold et al., 2011; Adachi et al., 2013). In our study, we explore the biological effects of diverse E-P associations with *Sox2* gene. This is an alternative way offering that scientist could study the transcription control of crucial TF genes, such as *Sox2*, without deletion of coding regions. With three E-P association KO cells, we focus on analyzing their DEGs and biological effects in mESCs. Our results reveal that these E-P associations involve in cooperative regulation of *Sox2* transcription. This may represent a quantitative way to explore gene transcription control. Thus, our study provides a potential new strategy for gene function study.

In GO analysis, cardiac function is enriched in RNA-Seq datasets across three KO cells. With further clustering, the list of genes (Table S5), which are all well annotated as cardiac function related, are further divided into three different subdivisions (Figure 6A). And their expression patterns are quite different among these three cells. For instance, the up-regulated DEGs mainly affect angiogenesis in E1-KO cell which have a little change in E2-KO and E3-KO cells (Figure 6A). Similar phenomenon is identified in metabolism related genes (Figure 6B) (Table S6). These imply that the same and well annotated gene function could show different expression patterns with disrupting different E-P association. As known, different gene expression patterns are closely correlated with its function(Hoffmann et al., 2002; Hagey et al., 2018). Thus, our finding might imply that the function of these genes needs more precisely exploration with their E-P associations.

The *Sox2* gene is highly expressed and interacting with multiple cofactors or E-P associations in mESCs (Figure 7). Its function heavily depends on different E-P associations. In our results, specific interaction (E1) regulates cell cycle self-renewal and cardiac function which are verified *in vitro* and *in vivo.* Indirect one (E2) affects metabolism and other aspect in cell function such as nervous system function which suggests it might potentially involve in *Sox2* transcription control through other mechanism. Common one (E3) involves in ion transport and metabolism. Additionally, it has significant influence on pluripotency of stem cells which downregulates genes such as *Klf4, Akt1, DPP3* and *TCL1* (Figure 7). These imply that common and indirect E-P associations might have some bypass pathways, which also indicates the molecular mechanism of *Sox2* gene expression and transcription control might be more complicated than previously anticipated.

In summary, diverse E-P associations are prevalent in gene transcription process. Our study indicates complicated E-P associations can cooperatively regulate *Sox2* function in various patterns. This offers an alternative way to explore gene expression and transcription control in spatiotemporal manner. In the future, E-P associations might be an indispensable process to comprehensively decipher gene function.

## Materials and Methods

### Cell culture

The mESCs cell line (E14, obtained from the American Type Culture Collection) is cultured on 0.1% gelatin-coated plates in ES medium (Knockout-DMEM containing 15% FBS, 100 U/mL Penicillin, 100 ug/mL Streptomycin, 0.1 mM MEM nonessential amino acids, 1 mM sodium pyruvate, 2 mM GlutaMAX, 0.1 mM 2-mercaptoethanol, 1000 U/mL LIF, 3 mM CHIR99021 [GSK3b inhibitor, Selleck], 1 mM PD0325901 [MEK inhibitor, Selleck]) and maintained in a pluripotent state in the absence of a feeder layer(Miyagi et al., 2006; Ying et al., 2008). Human embryonic kidney 293T cell (obtained from the American Type Culture Collection) are cultured in DMEM containing 10% FBS, 100 U/mL Penicillin, 100 ug/mL Streptomycin and 2 mM GlutaMAX.

### Enhancer validation and deletion

The activities of E1, E2, E3 are assayed using fluorescence indication assay. The vector is based on the modified pGL4.23, named pGL4.23-mCherry (PGL4.23 is digested by XbaI/NcoI to remove the original luc2 reporter gene CDS and replace it with mCherry.). Briefly, PCR-amplified E1 and E3 are inserted downstream from the mCherry gene by SalI/BamHI in the pGL4.23-mCherry vector and transfected into 293T cells on 12-well plates. E2 is inserted upstream from mCherry gene by KpnI/XhoI. All the transfections are carried out in the same manner. Briefly, 1 μg of DNA were transfected in mESCs or 293T cells using Fugene HD according to the manufacturer’s protocols.

To assay the role of enhancers in *Sox2* gene expression, we delete these regions in mouse ES cells using the CRISPR/Cas9 system. U6 promoter-driven gRNA cloning vector pGL3-U6-gRNA-PGK-puromycin (51133) and Cas9 expressing plasmid (44758) are purchased from Addgene (Cambridge, USA). Two pairs of gRNAs are designed according to the upstream and downstream of each enhancer (Table S1). All the four plasmids encoding gRNAs are assembled using the gRNA empty vector and co-transfected with Cas9 into E14 cells using Nucleofector II (Amaxa, Germany). The dosage of plasmids is 5 μg for 1 million cells (The proportion of gRNA and Cas9 plasmid is 2 to 1). Targeted clones are screened by PCR to identify deletions (primers in Table S1), and deletions are confirmed by sanger sequencing. Gene expression (normalized to *Gapdh*) is quantified by Quantitative real-time PCR (qPCR). Total RNA is extracted with Trizol (Invitrogen). Samples are treated with DNase I before reverse transcription using random priming and Superscript Reverse Transcriptase (Thermo Scientific), according to the manufacturer’s protocols. QPCR is performed using ABI Step one (Applied Biosystems) and Powerup SYBR Green Master Mix (Applied Biosystems). Gene expression is evaluated using the ΔΔCt method. The primers are showed in supplementary data (Table S1).

### RNA-seq analysis

RNA is isolated using Trizol (Invitrogen) from WT and enhancer KO ES cells. The 8 libraries (two biological duplicates in four groups) are constructed by VAHTSTM Total RNA-seq (H/M/R) Library Prep Kit (Vazyme Biotech, China) prior to multiplexed massively parallel sequencing (paired-end 150 base pairs [bp]) using the Illumina platform. Sequence data are submitted to the Gene Expression Omnibus (GEO) repository (GSE115750).

To ensure the quality of data used in further analysis, rRNA sequences are filtered out by SortMeRNA v2.0(Kopylova et al., 2012) (default parameter) firstly. Clean reads are obtained from the raw data by removing adapter-containing reads, poly-N-containing reads and low-quality reads using Trimmomatic (version 0.36)(Bolger et al., 2014). All downstream analyses are based on high quality clean data. Clean data is mapped to NCBI37/mm9 reference genome using TopHat v2.0.12(Trapnell et al., 2009). The parameter --read-mismatches and --library-type are set to 5 and fr-firststrand, respectively.

Differential expression analysis of enhancer-knock-out and WT mESCs groups is performed by Cuffdiff (2.2.1)(Trapnell et al., 2012). In this study, genes with q-value <0.05 and | log2 (fold change) | > 1 between the enhancer-knock-out and WT mESCs groups are identified as DEGs. GO and KEGG enrichment analyses of the DEGs are performed using DAVID 6.8(Huang da et al., 2009b; Huang da et al., 2009a) and KOBAS 3.0(Wu et al., 2006; Xie et al., 2011) software, respectively. Terms with p<0.05 and pathway with corrected *p*-value <0.05 are regarded as significant enrichment.

### Cell cycle and proliferation analysis

PI/RNase Staining Buffer is used for cell cycle assay. Cell sample is fixed and permeabilized with ice cold anhydrous ethanol. Then 0.5 mL buffer (for 1 million cells) is used for incubating for 15 minutes at room temperature before analysis. Fluorescence signal is detected using Beckman Coulter CytoFLEX flow cytometry system, and the cell cycle curve is fitted by ModFit software. The reagent kit for Edu cell proliferation assay is purchased from Guangzhou RiboBio Co., LTD, Guangdong, China. The mESCs proliferation assay is performed according to the kit instructions.

### Clonogenic assay

Cells are seeded at a density of 200 per 6-cm dish. After 5-7 days, the colonies are fixed with 4% paraformaldehyde at room temperature for 15 minutes and stained using an BCIP/NBT alkaline phosphatase (ALP) staining kit (Beyotime Biotechnology, China) according to a standard protocol. Clones that are larger than 100 microns in diameter are enumerated for each group.

### Neural differentiation assay

Neural differentiation of mESCs is performed with NDiff227 medium (Takara) in adherent monolayer culture conditions as described in Ying QL, et al(Ying et al., 2003). Plate feeder independent mESCs in Ndiff227 medium onto gelatin-coated tissue culture plastic and change medium every 1-2 days.

### Zebrafish Validation

The Zebrafish Validation of enhancer activity is referred to our previous work (Zhang et al., 2013)

## Supplementary Data

Supplementary Data are available online.

## Author contributions

Y.Z. conducts and designs the experiments. L.H. and S.K. performs the experiments. Q.L. and Q.H. conducts the bioinformatics work. X.Z. and Y.P. participates in discussions and give good advice. The manuscript is written by L.H., Q.L. and Y.Z. All authors read and approve the final manuscript.

## Acknowledgements

We thank Y. Cheng for discussion and advice on design of gRNA.

## Funding from

Agricultural Science and Technology Innovation Program of Chinese Academy of Agricultural Sciences. Guangdong Natural Science Foundation [2018A0303130009].

## Data availability

Sequence data are submitted to the Gene Expression Omnibus (GEO) repository (GSE115750).

## References

Adachi, K., Nikaido, I., Ohta, H., Ohtsuka, S., Ura, H., Kadota, M., Wakayama, T., Ueda, H. R. and Niwa, H. (2013) ‘Context-dependent wiring of Sox2 regulatory networks for self-renewal of embryonic and trophoblast stem cells’, Mol Cell 52(3): 380–92.

Ang, Y. S., Tsai, S. Y., Lee, D. F., Monk, J., Su, J., Ratnakumar, K., Ding, J., Ge, Y., Darr, H., Chang, B. et al. (2011) Wdr5 mediates self-renewal and reprogramming via the embryonic stem cell core transcriptional network’, Cell 145(2): 183–97.

Arnold, K., Sarkar, A., Yram, M. A., Polo, J. M., Bronson, R., Sengupta, S., Seandel, M., Geijsen, N. and Hochedlinger, K. (2011) ‘Sox2(+) adult stem and progenitor cells are important for tissue regeneration and survival of mice’, Cell Stem Cell 9(4): 317–29.

Avilion, A. A., Nicolis, S. K., Pevny, L. H., Perez, L., Vivian, N. and Lovell-Badge, R. (2003) ‘Multipotent cell lineages in early mouse development depend on SOX2 function’, Genes Dev 17(1): 126–40.

Bolger, A. M., Lohse, M. and Usadel, B. (2014) ‘Trimmomatic: a flexible trimmer for Illumina sequence data’, Bioinformatics 30(15): 2114–20.

DeMare, L. E., Leng, J., Cotney, J., Reilly, S. K., Yin, J., Sarro, R. and Noonan, J. P. (2013) ‘The genomic landscape of cohesin-associated chromatin interactions’, Genome Res 23(8): 1224–34.

Dowen, J. M., Fan, Z. P., Hnisz, D., Ren, G., Abraham, B. J., Zhang, L. N., Weintraub, A. S., Schujiers, J., Lee, T. I., Zhao, K. et al. (2014) ‘Control of cell identity genes occurs in insulated neighborhoods in mammalian chromosomes’, Cell 159(2): 374–387.

Drissen, R., Guyot, B., Zhang, L., Atzberger, A., Sloane-Stanley, J., Wood, B., Porcher, C. and Vyas, P. (2010) ‘Lineage-specific combinatorial action of enhancers regulates mouse erythroid Gata1 expression’, Blood 115(17): 3463–71.

Hagey, D. W., Klum, S., Kurtsdotter, I., Zaouter, C., Topcic, D., Andersson, O., Bergsland, M. and Muhr, J. (2018) ‘SOX2 regulates common and specific stem cell features in the CNS and endoderm derived organs’, PLoS Genet 14(2): e1007224.

Hnisz, D., Shrinivas, K., Young, R. A., Chakraborty, A. K. and Sharp, P. A. (2017) ‘A Phase Separation Model for Transcriptional Control’, Cell 169(1): 13–23.

Hoffmann, R., Seidl, T., Neeb, M., Rolink, A. and Melchers, F. (2002) ‘Changes in gene expression profiles in developing B cells of murine bone marrow’, Genome Res 12(1): 98–111.

Huang da, W., Sherman, B. T. and Lempicki, R. A. (2009a) ‘Bioinformatics enrichment tools: paths toward the comprehensive functional analysis of large gene lists’, Nucleic Acids Res 37(1): 1–13.

Huang da, W., Sherman, B. T. and Lempicki, R. A. (2009b) ‘Systematic and integrative analysis of large gene lists using DAVID bioinformatics resources’, Nat Protoc 4(1): 44–57.

Jager, R., Migliorini, G., Henrion, M., Kandaswamy, R., Speedy, H. E., Heindl, A., Whiffin, N., Carnicer, M. J., Broome, L., Dryden, N. et al. (2015) ‘Capture Hi-C identifies the chromatin interactome of colorectal cancer risk loci’, Nat Commun 6: 6178.

Ji, X., Dadon, D. B., Powell, B. E., Fan, Z. P., Borges-Rivera, D., Shachar, S., Weintraub, A. S., Hnisz, D., Pegoraro, G., Lee, T. I. et al. (2016) ‘3D Chromosome Regulatory Landscape of Human Pluripotent Cells’, Cell Stem Cell 18(2): 262–75.

Kieffer-Kwon, K. R., Tang, Z., Mathe, E., Qian, J., Sung, M. H., Li, G., Resch, W., Baek, S., Pruett, N., Grontved, L. et al. (2013) ‘Interactome maps of mouse gene regulatory domains reveal basic principles of transcriptional regulation’, Cell 155(7): 1507–20.

Kleinjan, D. A. and van Heyningen, V. (2005) ‘Long-range control of gene expression: emerging mechanisms and disruption in disease’, Am J Hum Genet 76(1): 8–32.

Kopylova, E., Noe, L. and Touzet, H. (2012) ‘SortMeRNA: fast and accurate filtering of ribosomal RNAs in metatranscriptomic data’, Bioinformatics 28(24): 3211–7.

Krijger, P. H. and de Laat, W. (2016) ‘Regulation of disease-associated gene expression in the 3D genome’, Nat Rev Mol Cell Biol 17(12): 771–782.

Li, G., Ruan, X., Auerbach, R. K., Sandhu, K. S., Zheng, M., Wang, P., Poh, H. M., Goh, Y., Lim, J., Zhang, J. et al. (2012a) ‘Extensive promoter-centered chromatin interactions provide a topological basis for transcription regulation’, Cell 148(1–2): 84–98.

Li, H., Collado, M., Villasante, A., Matheu, A., Lynch, C. J., Canamero, M., Rizzoti, K., Carneiro, C., Martinez, G., Vidal, A. et al. (2012b) ‘p27(Kip1) directly represses Sox2 during embryonic stem cell differentiation’, Cell Stem Cell 11(6): 845–52.

Li, Y., Rivera, C. M., Ishii, H., Jin, F., Selvaraj, S., Lee, A. Y., Dixon, J. R. and Ren, B. (2014) ‘CRISPR reveals a distal super-enhancer required for Sox2 expression in mouse embryonic stem cells’, PLoS One 9(12): e114485.

Liu, X., Zhang, Y., Chen, Y., Li, M., Zhou, F., Li, K., Cao, H., Ni, M., Liu, Y., Gu, Z. et al. (2017) ‘In Situ Capture of Chromatin Interactions by Biotinylated dCas9’, Cell 170(5): 1028–1043 e19.

Liu, Z., Legant, W. R., Chen, B. C., Li, L., Grimm, J. B., Lavis, L. D., Betzig, E. and Tjian, R. (2014) ‘3D imaging of Sox2 enhancer clusters in embryonic stem cells’, Elife 3: e04236.

Long, H. K., Prescott, S. L. and Wysocka, J. (2016) ‘Ever-Changing Landscapes: Transcriptional Enhancers in Development and Evolution’, Cell 167(5): 1170–1187.

Masui, S., Nakatake, Y., Toyooka, Y., Shimosato, D., Yagi, R., Takahashi, K., Okochi, H., Okuda, A., Matoba, R., Sharov, A. A. et al. (2007) ‘Pluripotency governed by Sox2 via regulation of Oct3/4 expression in mouse embryonic stem cells’, Nat Cell Biol 9(6): 625–35.

Miyagi, S., Nishimoto, M., Saito, T., Ninomiya, M., Sawamoto, K., Okano, H., Muramatsu, M., Oguro, H., Iwama, A. and Okuda, A. (2006) ‘The Sox2 regulatory region 2 functions as a neural stem cell-specific enhancer in the telencephalon’, J Biol Chem 281(19): 13374–81.

Narva, E., Rahkonen, N., Emani, M. R., Lund, R., Pursiheimo, J. P., Nasti, J., Autio, R., Rasool, O., Denessiouk, K., Lahdesmaki, H. et al. (2012) ‘RNA-binding protein L1TD1 interacts with LIN28 via RNA and is required for human embryonic stem cell self-renewal and cancer cell proliferation’, Stem Cells 30(3): 452–60.

Rowan, S., Siggers, T., Lachke, S. A., Yue, Y., Bulyk, M. L. and Maas, R. L. (2010) ‘Precise temporal control of the eye regulatory gene Pax6 via enhancer-binding site affinity’, Genes Dev 24(10): 980–5.

Rubin, A. J., Barajas, B. C., Furlan-Magaril, M., Lopez-Pajares, V., Mumbach, M. R., Howard, I., Kim, D. S., Boxer, L. D., Cairns, J., Spivakov, M. et al. (2017) ‘Lineage-specific dynamic and pre-established enhancer-promoter contacts cooperate in terminal differentiation’, Nat Genet 49(10): 1522–1528.

Rubtsov, M. A., Polikanov, Y. S., Bondarenko, V. A., Wang, Y. H. and Studitsky, V. M. (2006) ‘Chromatin structure can strongly facilitate enhancer action over a distance’, Proc Natl Acad Sci U S A 103(47): 17690–5.

Seitan, V. C., Faure, A. J., Zhan, Y., McCord, R. P., Lajoie, B. R., Ing-Simmons, E., Lenhard, B., Giorgetti, L., Heard, E., Fisher, A. G. et al. (2013) ‘Cohesin-based chromatin interactions enable regulated gene expression within preexisting architectural compartments’, Genome Res 23(12): 2066–77.

Sikorska, M., Sandhu, J. K., Deb-Rinker, P., Jezierski, A., Leblanc, J., Charlebois, C., Ribecco-Lutkiewicz, M., Bani-Yaghoub, M. and Walker, P. R. (2008) ‘Epigenetic modifications of SOX2 enhancers, SRR1 and SRR2, correlate with in vitro neural differentiation’, JNeurosci Res 86(8): 1680–93.

Trapnell, C., Pachter, L. and Salzberg, S. L. (2009) ‘TopHat: discovering splice junctions with RNA-Seq’, Bioinformatics 25(9): 1105–11.

Trapnell, C., Roberts, A., Goff, L., Pertea, G., Kim, D., Kelley, D. R., Pimentel, H., Salzberg, S. L., Rinn, J. L. and Pachter, L. (2012) ‘Differential gene and transcript expression analysis of RNA-seq experiments with TopHat and Cufflinks’, Nat Protoc 7(3): 562–78.

Tsai, P. F., Dell’Orso, S., Rodriguez, J., Vivanco, K. O., Ko, K. D., Jiang, K., Juan, A. H., Sarshad, A. A., Vian, L., Tran, M. et al. (2018) ‘A Muscle-Specific Enhancer RNA Mediates Cohesin Recruitment and Regulates Transcription In trans’, Mol Cell 71(1): 129–141.e8.

Weintraub, A. S., Li, C. H., Zamudio, A. V., Sigova, A. A., Hannett, N. M., Day, D. S., Abraham, B. J., Cohen, M. A., Nabet, B., Buckley, D. L. et al. (2017) ‘YY1 Is a Structural Regulator of Enhancer-Promoter Loops’, Cell 171(7): 1573–1588 e28.

Wu, J., Mao, X., Cai, T., Luo, J. and Wei, L. (2006) ‘KOBAS server: a web-based platform for automated annotation and pathway identification’, Nucleic Acids Res 34(Web Server issue): W720–4.

Xie, C., Mao, X., Huang, J., Ding, Y., Wu, J., Dong, S., Kong, L., Gao, G., Li, C. Y. and Wei, L. (2011) ‘KOBAS 2.0: a web server for annotation and identification of enriched pathways and diseases’, Nucleic Acids Res 39(Web Server issue): W316–22.

Yamamizu, K., Schlessinger, D. and Ko, M. S. (2014) ‘SOX9 accelerates ESC differentiation to three germ layer lineages by repressing SOX2 expression through P21 (WAF1/CIP1)’, Development 141(22): 4254–66.

Yanling Peng and Yubo Zhang (2018) ‘Enhancer and super □enhancer: Positive regulators in gene transcription’. Anim Model Exp Med 3(1):169–179.

Ying, Q. L., Nichols, J., Chambers, I. and Smith, A. (2003) ‘BMP induction of Id proteins suppresses differentiation and sustains embryonic stem cell self-renewal in collaboration with STAT3’, Cell 115(3): 281–92.

Ying, Q. L., Wray, J., Nichols, J., Batlle-Morera, L., Doble, B., Woodgett, J., Cohen, P. and Smith, A. (2008) ‘The ground state of embryonic stem cell self-renewal’, Nature 453(7194): 519–23.

Zhang, Y., Wong, C. H., Birnbaum, R. Y., Li, G., Favaro, R., Ngan, C. Y., Lim, J., Tai, E., Poh, H. M., Wong, E. et al. (2013) ‘Chromatin connectivity maps reveal dynamic promoter-enhancer long-range associations’, Nature 504(7479): 306–310.

Zhou, H. Y., Katsman, Y., Dhaliwal, N. K., Davidson, S., Macpherson, N. N., Sakthidevi, M., Collura, F. and Mitchell, J. A. (2014) ‘A Sox2 distal enhancer cluster regulates embryonic stem cell differentiation potential’, Genes Dev 28(24): 2699–711.

